# *In vitro* inhibition of *H. pylori* in a preferential manner using bioengineered *L. lactis* releasing guided Antimicrobial peptides

**DOI:** 10.1101/2021.06.11.448109

**Authors:** Ankan Choudhury, Patrick Ortiz, Christopher M. Kearney

## Abstract

**Objectives:** Targeted therapies seek to selectively eliminate a pathogen without disrupting the resident microbial community. This is even more important when a pathogen like *H. pylori* resides in stomach, a sensitive microbial ecosystem. Using a probiotic like *Lactococcus lactis* and bioengineering it to release a guided Antimicrobial Peptide (AMP) targeted towards the pathogen offers a pathway to specifically knock-out the deleterious species and not disturbing the stomach microbiome.

**Results:** Three AMPs, Alyteserin, CRAMP and Laterosporulin, were genetically fused to a guiding peptide MM1, which selectively binds to Vacuolating Toxin A (VacA) of *H. pylori* and cloned into an excretory vector pTKR inside *L. lactis*. When cultured together *in vitro*, the *L. lactis* bioengineered with guided AMPs selectively killed *H. pylori* when compared to *E. coli* or *Lactobacillus plantarum*, as determined by qPCR. Chemically synthesized Alyteserin and MM1-Alyteserin showed similar preferential inhibition of *H. pylori* when compared against *E. coli*, with the MIC of MM1-Alyteserin becoming significantly higher for *E. coli* than Alytserin whereas no such effet was observed against *H. pylori*.

**Conclusions:** Probiotics bioengineered to excrete guided AMPs can be a novel and useful approach for combating pathogens without endangering the natural microbial flora. Given the wealth of AMPs and guiding ligands, both natural and synthetic, this approach can be adapted to develop a diverse array of chimeric guided AMPs and can be cloned into probiotics to create a safe and effective alternative to conventional chemical antibiotics.

## Introduction

*Helicobacter pylori* is the source of one of the most prevalent infections in the world, with over 50% prevalence in many countries but often over 90% in Africa and East Asia (Salih, 2009). Over 60% of cases of gastric cancer can be attributed to *H. pylori* infection (Correa and Piazuelo, 2011), making it one of the most widespread cancers caused by an infectious agent (Wroblewski et al., 2010). Multidrug resistant strains of *H. pylori* constitute an increasing portion of *H. pylori* infections, from >10% in European countries to >40% of infections in Peru (Boyanova et al., 2019). Newest treatment regimens for *H. pylori* infection include triple and quadruple antibiotic therapies to match the growing challenge of antibiotic resistance. Such therapeutic regimens include combinations of amoxicillin, tetracycline, bismuth, metronidazole, clarithromycin, and more. In return, quadruple, quintuple, and sextuple antibiotic-resistant strains have been detected (Boyanova et al., 2019). This escalation of antibiotic resistance in *H. pylori* has heightened the need for new therapeutic strategies to combat infection. The multiple actions of these antibiotics such as rRNA inhibition, β-lactams, nucleic acid inhibitors also deleteriously effect off target bacteria, and a growing list of antibiotics administered to curb a single infection increases the dysbiosis of commensal microbiota caused by killing of off-target bacteria (Becattini et al., 2016; Langdon et al., 2016; Zarrinpar et al., 2018). Antibiotic-associated Dysbiosis often precipitates into intestinal inflammatory diseases like colitis (Strati et al., 2021), worsens neuro-immune mechanisms and viscerosensory functionalities (Aguilera et al., 2015) and often makes way for bloom of pathogens (Vangay et al., 2015) creating other possibly more serious infectious diseases. This presents a dilemma, as stronger small molecule antibiotics are required to kill bacteria with ever-evolving antibiotic resistance mechanisms, but stronger antibiotics kill a wider variety of commensal bacteria (Becattini et al., 2016; Langdon et al., 2016; Zarrinpar et al., 2018).

To meet the challenges associated with this infection, one strategy proposed has been the use of AMPs. AMP refers to a broad group of short, usually cationic peptides with bactericidal or bacteriostatic properties (Lei et al., 2019). Because many of them exhibit a broad mechanism of action that forms pores in bacterial membranes, it has been suggested that it may be more difficult for bacteria to evolve resistance mechanisms to these peptides than traditional antibiotic drugs, though resistance can still occur (Assoni et al., 2020; Di et al., 2020; El Shazely et al., 2020). While the broad category of AMP comprises many diverse peptides that exhibit some antimicrobial activity, several specific types of AMP have been demonstrated to effectively kill *H. pylori*. Cathelicidins such as LL-37 and its murine homolog Cathelin-related Antimicrobial Peptide (CRAMP) have been demonstrated to effectively kill *Helicobacter pylori* in both *in vitro* and *in vivo* experiments (Hase et al., 2003; Zhang et al., 2016, 2013). Bacteriocins are small, stable AMPs released by other bacteria, that have broad bactericidal ability against a variety of gram-positive and gram-negative bacteria including *H. pylori* (Neshani et al., 2019). Among them, Type IId bacteriocins including Laterosporulin has been well documented for their bactericidal activity with well-established mechanisms (Baindara et al., 2016; Singh et al., 2015). As more novel AMPs are discovered, a catalog of AMPs with activity against *H. pylori* has grown, showing promise as potential therapeutics.

While many of these AMPs have demonstrated effective antibacterial activity towards *H. pylori*, they also kill many other bacterial taxa. The double-edged sword of antibacterial therapies is that they have the unintended consequence of killing commensal microbiota. To deal with the problem of off target killing there have been several proposed solutions. Some AMPs naturally have increased activity towards specific bacterial taxa, and if utilized properly might avoid causing dysbiosis of microbiota in certain settings. Another option has been to modify AMPs, making chimeric peptides that use a short glycine linker and a guide peptide to “target” a specific taxon. Such guided antimicrobial peptides (gAMPs) have been shown to be effective in several settings against a variety of bacteria (Choudhury et al., 2020; Eckert et al., 2012, 2006; Kim et al., 2020). In some cases, such constructs can be made to increase the toxicity of a relatively weak AMP towards a targeted bacterium (Eckert et al., 2006), whereas in others it has been demonstrated to decrease toxicity of a potent AMP towards off-target bacteria (Choudhury et al., 2020). Furthermore, while studies have shown the bactericidal effects of such gAMPs in an *in-vivo* setting, the selectivity of these constructs has not been demonstrated *in-vivo* to ascertain if the microbiota are relatively undisturbed; nor has a gAMP been utilized against *H. pylori*. One of the reasons for this is that delivery of engineered peptides may be difficult. Intraperitoneal injections of purified peptide have been used for gut infections, but infections of the stomach require a delivery mechanism that will stand up to low pH conditions, peptidases, and provide delivery at the site of the infection. Antimicrobial peptides, being proteinaceous, are at a greater risk of enzymatic degradation through oral routes (Moncla et al., 2011; Svenson et al., 2008) and the high gastric acidity and peptidolytic enzymes cause breakdown of proteins and peptides when ingested orally. To avoid this gastric degradation, drugs are often delivered through systemic injection. For peptides, this is problematic as the size and high molecular weight of proteinaceous drug make it an easier target for opsonization and neutralization by the blood complement system (Vaucher et al., 2011). Thus, for having the desired therapeutic effect the peptidic drug will have to survive the degradation in gut and reach the site of action. Encasing the antimicrobial peptide is in a delivery system that masks it to survive the journey in the oral delivery and release it once the site is reached would be of great help and would help in microbial infections along the gut for which oral delivery of drugs is necessary.

Employing food grade bacterial systems like the lactic acid bacteria can solve the problem of the peptide’s survival through degradative environments such as the gastrointestinal tract (Steidler et al., 2003). These bacteria are adapted to survive, propagate and produce and secrete their indigenous proteins in low pH conditions of the stomach. Encoding the chimeric antimicrobial peptide into a secretion vector inside such lactic acid bacteria will ensure that the protein will survive the journey into the gastrointestinal tract and be released from the cell into the site of infection (Jeong et al., 2006; Li et al., 2011). The cells will act as a sustained release platform as the expression of the protein will happen over a time. The cells will also replicate and maintain a colony of drug-releasing bacteria for an extended period (Drouault et al., 1999), unlike conventional drug delivery system. This reduces the number of dosages required to maintain the effective drug level for treatment of the infection. The vector can also be modified to contain an inducible promoter that is pH dependent (de Vos, 1999; Madsen et al., 1999), like the heat shock and nitrogen dependent constitutive promoters. A promoter that is induced by low acidic pH, like P1, P2 and P170 (de Vos, 1999; Madsen et al., 2005, 1999), will enable the lactic acid bacteria to express and secrete the encoded peptide only when it is exposed to such conditions at the target location in the gastric system. Thus, a lactic acid bacterium containing a secretion vector with a pH inducible promoter driving AMP expression constitutes an excellent sustained release drug delivery system that will protect the peptidic drug from the enzymatic degradation in the gastrointestinal tract and deliver it to the proper target site. Lactic acid bacteria in the genera *Lactobacillus* and *Lactococcus* have been a part of the human diet for millennia. There have been numerous strains of lactic acid bacteria that are considered safe to consume and graded by the FDA as such (Nutrition, 2020) Since early 2000s, *Lactococcus lactis* has proven to be an excellent method for delivering engineered peptides *in-situ* from what has since burgeoned a variety of engineered probiotic bacteria. These engineered probiotics have been used to deliver signaling peptides, dyes, interleukins, and even unmodified AMPs (Foligne et al., 2007; Steidler et al., 2003, 1998). *Lactococcus lactis* also holds the distinction of being the first genetically engineered organism to be used for a therapeutic application in humans (Braat et al., 2006).

In this paper, we will demonstrate the efficacy of *L. lactis* bioengineered to secrete AMPs and gAMPs to inhibit *H. pylori* when co-cultured *in vitro*. We selected a suitable vector for such purpose and then cloned it with 3 AMPs of our choice and their corresponding gAMPs. The gAMPs were created by fusing a guiding peptide to the N-terminus of the corresponding AMP. The principle behind our gAMPS was to find a unique and attractive target protein on the surface of *H. pylori*, for which we picked Vacuolating cytotoxin A (VacA). This protein expressed by *H. pylori* is a major cause for its pathogenicity and can either be released into extracellular space (Cover and Blaser, 1992; Foegeding et al., 2016; Snider et al., 2016) or a significant portion of it is left in the bacterial cell surface (Foegeding et al., 2016; McClain and Cover, 2006; Telford et al., 1994; Voss et al., 2014) which is then transferred to host cell via contact dependent mechanisms (Ilver et al., 2004; Keenan et al., 2000; McClain and Cover, 2006). VacA toxin (comprised of the p33 and p55 subunits) binds to host cells and is internalized, causing severe “vacuolation” characterized by the accumulation of large vesicles that possess hallmarks of both late endosomes and early lysosomes (Foegeding et al., 2016; Palframan et al., 2012). The development of “vacuoles” has been attributed to the formation of VacA anion-selective channels in membranes. Apart from its vacuolating effects, it has recently become clear that VacA also directly affects mitochondrial function (Foo et al., 2010). The VacA toxin binds to stomach lining cells by associating with the lipid rafts on the cell membrane which causes it to be internalized by the cell and promote vacuole formation (Fiocca et al., 1999). Once internalized, the p34 subunit of VacA toxin also forms an anionic pore into the mitochondrial membrane and interfere with its function (Domańska et al., 2010). The vacA toxin has been studied for being a major virulence factor and hence its structures and associations are also explored. One such protein whose association with the VacA toxin was explored was Multimerin-1(Satoh et al., 2013). Multimerin-1 is a protein expressed on the surface of human platelets and is bound by the VacA toxin when it induces platelet CD62P expression76. It was shown that VacA would bind with a specified region of multimerin-1 from AA 321-340 stronger than any other region, making it the purported region for attachment with the platelets (Satoh et al., 2013). We used this peptide sequence, fused to the AMP, as a targeting domain against *H. pylori*. Expression will be driven by the low pH induced secretion vector in *L. lactis*. This will yield a live vector delivery system for AMP targeted towards only *H. pylori* and sparing other bacteria.

For expressing the AMPs and gAMPs in *L. lactis* we picked the vector pT1NX (Steidler et al., 2004, 1998; van Asseldonk et al., 1990; Waterfield et al., 1995). pT1NX contains a native *L. lactis* promoter called P1 (Madsen et al., 2005) which is induced by the low pH conditions like in the stomach. P1 is also induced by thermal stress and have been seen to increase expression of proteins 500-fold when growth conditions are changed from 24°C to around 40°C, which is closer to a human physiological condition (de Vos, 1999). pT1NX also contains the native signal peptide usp45 (van Asseldonk et al., 1990) that secretes any expressed protein placed downstream of it in the ORF. Thus, placing the AMP or the gAMP placed downstream in the ORF of the vector will ensure that the usp45 signal peptide will secrete it outside the *L. lactis* cell surface and this expression and subsequent secretion will be governed by the P1 promoter which will trigger expression only in low pH conditions. The pT1NX vector was further modified by adding a Kanamycin Resistant domain to make it a dual bacteria vector and making it possible to be cloned into *E. coli* along with *L. lactis*.

## Methods and Materials

### Cloning Antimicrobial peptides(AMPs) and guided Antimicrobial peptides (gAMPs) in L. lactis

The ORFs of the AMPs (Table 3.1), codon-optimized for *Lactococcus lactis* were cloned into the modified pT1NX plasmid (pT1NX-Kanamycin resistant/pTKR, Figure 3.1) in between the restriction enzyme sites BamHI and SpeI by replacing the spaX protein of the original plasmid. The P1 promoter upstream of the BamHI cut-site controls the downstream expression as a constitutive promoter which is upregulated by low pH. The usp45 gene immediately upstream of BamHI site is an endogenous signal peptide of *Lactococcus* species that transports the attached protein to extracellular location.

After ligation of the AMP/gAMP into pTKR vector, it was transformed into *E. coli* (10β, NEB) and plated onto Kanamycin selective plate. The pT1NX plasmid (LMBP 3498) has erythromycin resistance but pTKR is a dual vector with kanamyicin resistance for cloning into electrocompetent *E. coli* (10β, NEB) for plasmid propagation. Extracted plasmid from the *E. coli* was then electroporated into electrocompetent *L. lactis* MG1363 (LMBP 3019) and plated on erythromycin selective GM17 plate (30°C, microaerobic, overnight). After screening for the presence of the AMP/gAMP ORFs with PCR, selected colonies are propagated in liquid cultures of M17 broth with Glucose (0.5% w/v) in the presence of erythromycin (5 µg/ml).

### *In vitro* assay: Co-culture assays with cloned *L. lactis* and *H. pylori* SS1

Desired variant of *L. lactis* cloned with AMP/tAMP were propagated from glycerol stocks and grown in GM17 broth overnight with erythromycin (5 µg/ml) with no shaking. *H. pylori* SS1 stocks were first propagated on Blood-Tryptic Soy agar (TSA) overnight with microaerobic condition and >5% CO2 environment. Then colonies from the plate were transferred to a TS broth with 5% Newborn Calf serum (NCS) and grown overnight under microaerobic condition and >5% CO2 environment. The *L. lactis* cultures were serially diluted in a 96 well culture plate with TS broth to make up a volume of 100 µL. To each well, 10 µL of the overnight *H. pylori* culture were added and each well volume was brought up to 200 µL with more TS broth. The plate is left to grow overnight in a microaerobic environment with >5% CO_2_. After 24 hrs, the each well content from the culture plate is transferred on a 96 well PCR plate. That PCR plate was sealed and heated for 15 min at 100°C and chilled at 4°C for 5 min. Then the plate was centrifuged at 2000g for 2 min and the supernatant was used as the template for qPCR. The qPCR was done using primers for VacA gene to quantify *H. pylori* (forward: 5’-ATGGAAATACAACAAACACAC-3’, reverse: 5’-CTGCTTGAATGCGCCAAAC-3’) and primers for *acma* gene for quantifying *L. lactis* (forward: 5’ GGAGCTCGTGAAAGCTGACT 3’, reverse: 5’ GCCGGAACATTGACAACCAC 3’). Standard curves for *H. pylori* and *L. lactis* were constructed by determining CT values for different dilutions of the overnight cultures of the respective bacteria (1/10, 1/100, 1/1000, 1/10000) in the qPCR plates, the CFUs for the dilutions were determined by plating on their respective agar plates. Same procedure was followed by co-culturing *Lactobacillus plantarum* and *E. coli* with serially diluted cultures of *L. lactis* for 24 hrs and their amount was determined by qPCR using primers for species specific genes for either bacteria. The amount of *L. lactis* added to the co-cultures of all the three assays ranged from around 4000 to 512000 CFU/ul.

### In vitro assay of chemically synthesized AMP and gAMP against H. pylori and E. coli

The peptide Alyteserin (2.28 kD) and its guided counterpart MM1-Alyteserin (5.02 kD) were synthesized and purchased from GenScript USA. Aliquots of the AMP/gAMP were used to plate 96 well bactericidal assay using both *H. pylori* and *E. coli*. AMP/ gAMP were serially diluted to achieve a range of concentrations from 800 µM to 6.25µM in each wells. Each well had a total volume of 100 µl with 50 µl of peptide solution in PBS, 45 µl of growth media (TSB with 5% NCS for *H. pylori* and Lennox Broth for *E. coli*) and 5 µl of bacterial culture grown overnight and diluted to OD600 of 0.6. The assay plates for *H. pylori* were grown in microaerobic condition with >5% CO2 at 37°C while *E. coli* plates were cultured in aerobic conditions overnight at 37°C. The MICs were determined by the concentration of the last well in ascending order that had bacterial growth associated turbidity. The inhibition of bacterial growth in the clear wells were confirmed by plating the clear well contents on a Mueller-Hinton agar at respective microaerobic and aerobic conditions.

## Results

### L. lactis cloned with gAMPs selectively killed H. pylori when co-cultured in vitro

On culturing together for 24 hrs, the cloned *L. lactis* showed a pronounced bactericidal activity towards *H. pylori* as determined by the qPCR method (Figure 2). The value of the *H. pylori* titer were calculated by using a concentration curve with Ct values found for CFU/ul values of *H. pylori* when plated. The gAMP containing *L. lactis* had similar bactericidal effect on *H. pylori* as their corresponding AMPs, except for MM1-Alyteserin showing a significant reduction of *H. pylori* CFUs at *L. lactis* doses around 64,000 CFU/ul or higher. The gAMP-*L. lactis*, at a concentration of 64,000 CFU/ul of *L. lactis*, reduced the *H. pylori* load to 1/30^th^ of the *H. pylori* load when co-cultured with *L. lactis* with empty vector at similar concentration on average, with MM1-Alyteserin-*L. lactis* reducing it to 1/1000^th^. The average reduction of *H. pylori* by the AMP-*L. lactis* was only 1/6^th^ of that with the *L. lactis* containing empty vector, with Laterosporulin-*L. lactis* performing the best at 1/60^th^ reduction at 64,000 CFU/ul. The reduction of *H. pylori* load increased with the concentration of *L. lactis* for both AMP and gAMP containing *L. lactis*.

**FIGURE 1.**
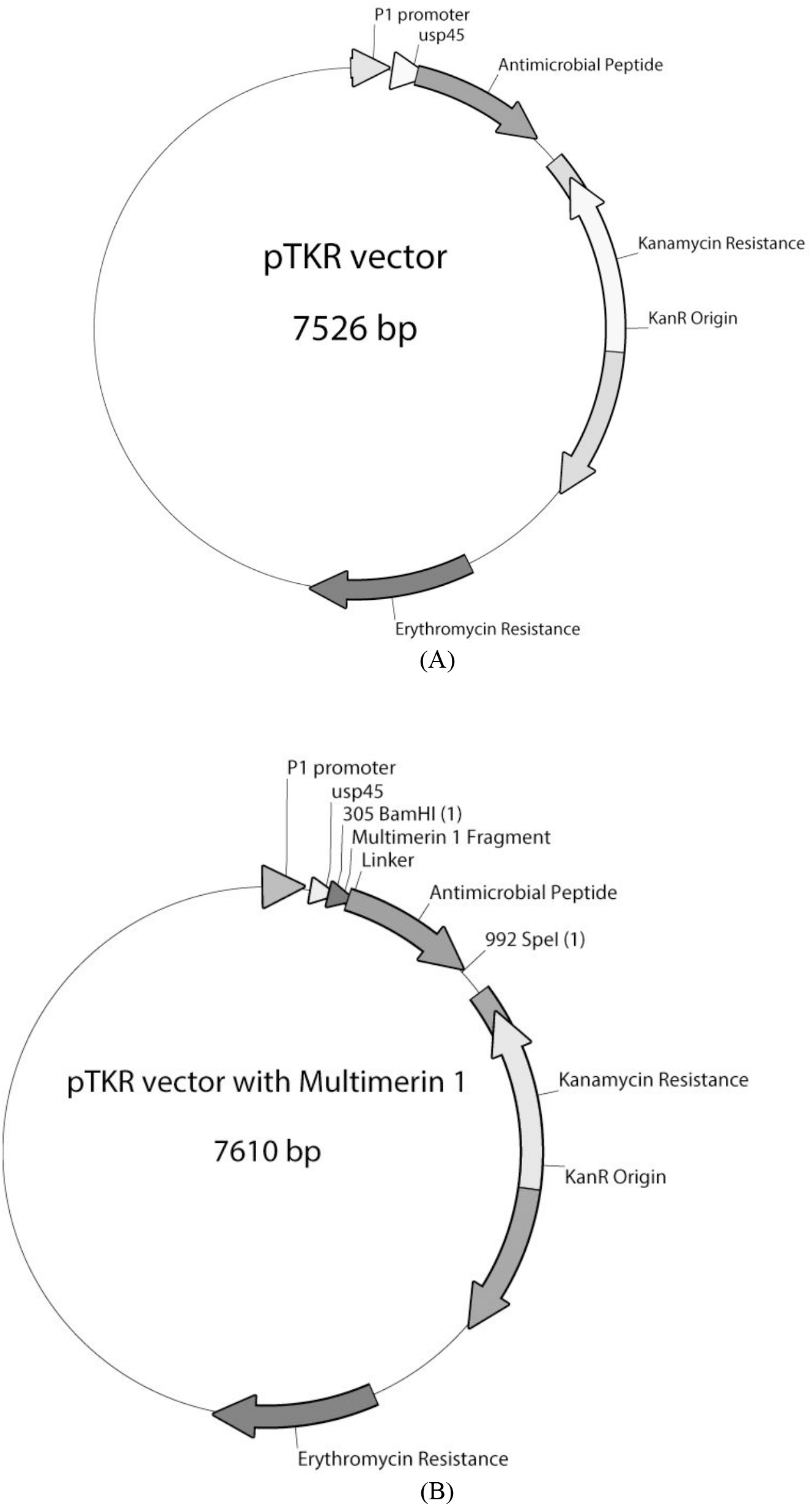
pTKR (pT1NX with kanamycin resistance gene) vector with the AMP sequence (A) and the MM1-AMP/ gAMP sequence (B) in between the BamHI and SpeI cutsites downstream of P1 promoter and usp45 sequence.

**FIGURE 2.**
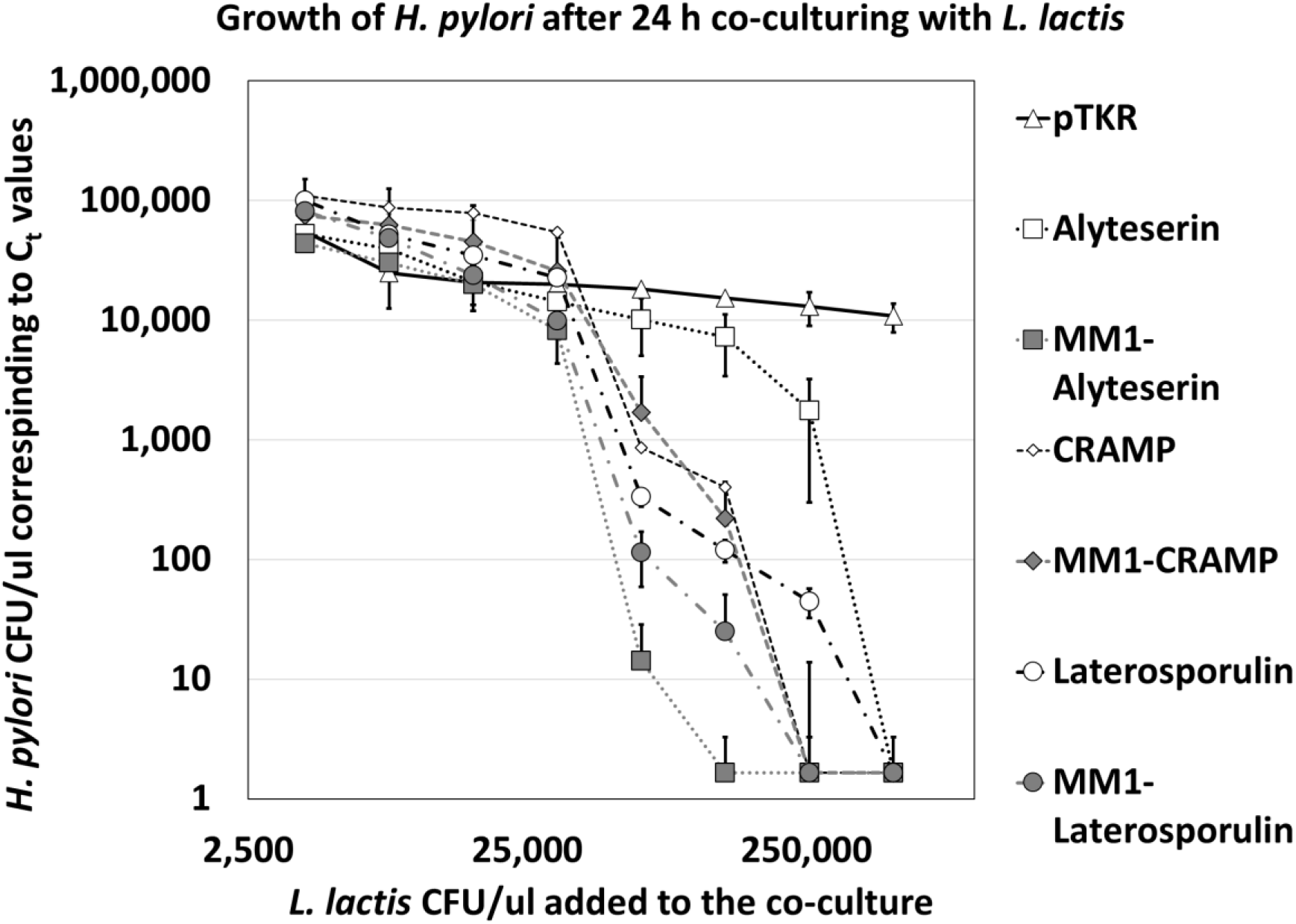
The CFU/µl of *H. pylori* grown in the co-culture after 24 hrs of incubation, estimated from the corresponding C_t_ values determined by qPCR, plotted against the CFU/µl of *L. lactis* added to the co-culture

But when co-cultured with *L. plantarum* (Figure 3) and *E. coli* (Figure 4), the gAMP-*L. lactis* had significantly less bactericidal action on either bacterium compared to their corresponding AMP-*L. lactis*. The difference in reduction of both *E. coli* and *L. plantarum* was evident even at the lowest concentration of *L. lactis* given (4000 CFU/µl). The bacterial load of *E. coli* and *L. plantarum* for the gAMP-*L. lactis* by 3.4x and 1.8x times higher on average respectively than the bacterial load when cultured with AMP-*L. lactis* at even 4000 CFU/µl, and the difference tapered with the increase in load of *L. lactis* upto 512000 CFU/µl. Hence, we can see that the *L. lactis* releasing gAMPs had significantly lesser killing potential for non-*H. pylori* bacteria than the *L. lactis* releasing corresponding AMPs, and this difference is only erased with exponentially higher doses of *L. lactis*. But inversely, gAMP releasing *L. lactis* killed *H. pylori* at almost similar rate with the AMP-*L. lactis* with MM1-Alyteserin performing the best among all the 6 AMP and gAMP tested.

**FIGURE 3.**
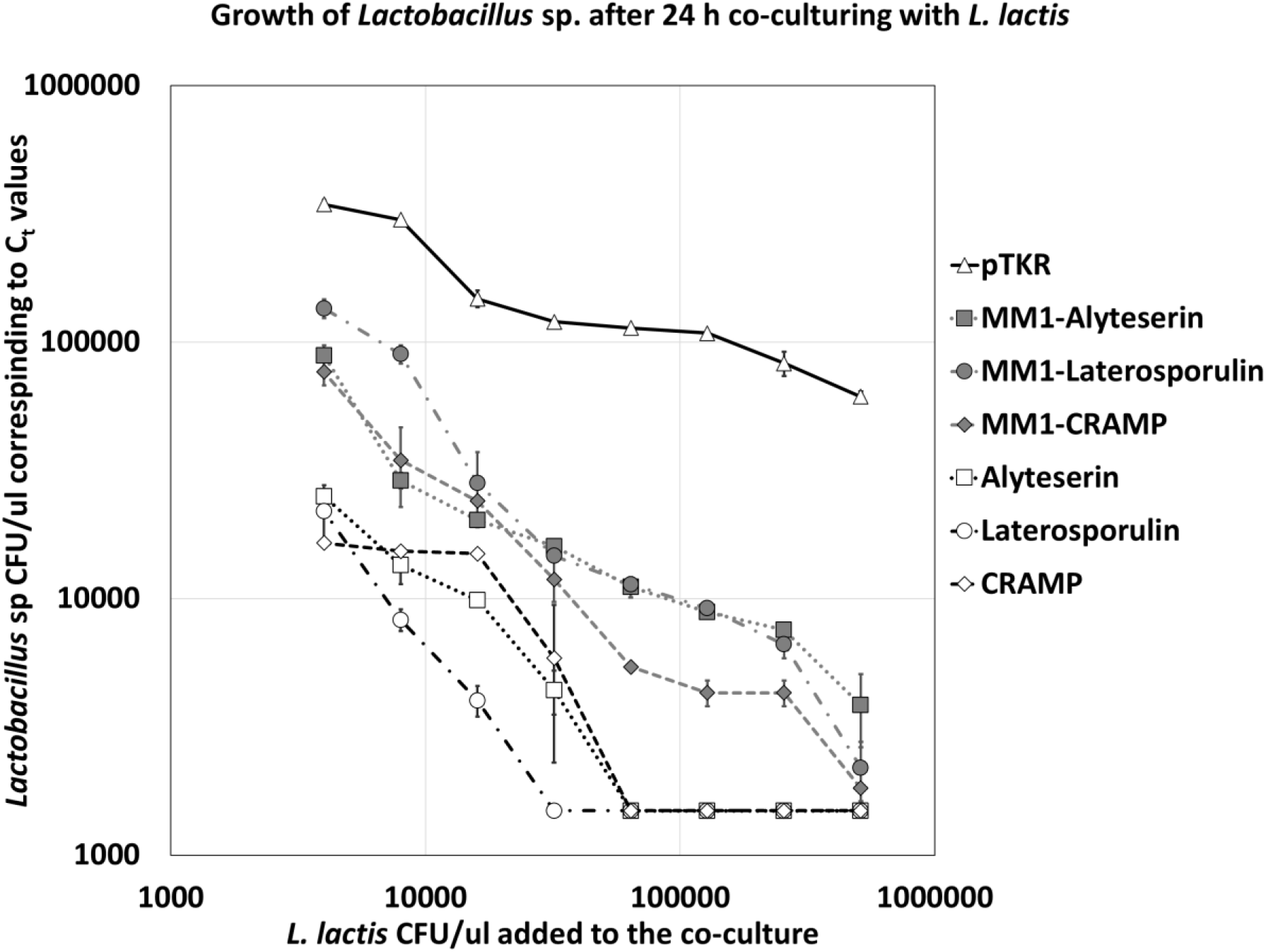
The CFU/µl of *L. plantarum* found in the co-culture after 24 hrs of incubation, estimated from the corresponding C_t_ values determined by qPCR, plotted against the CFU/µl of *L. lactis* added to the co-culture

**FIGURE 4.**
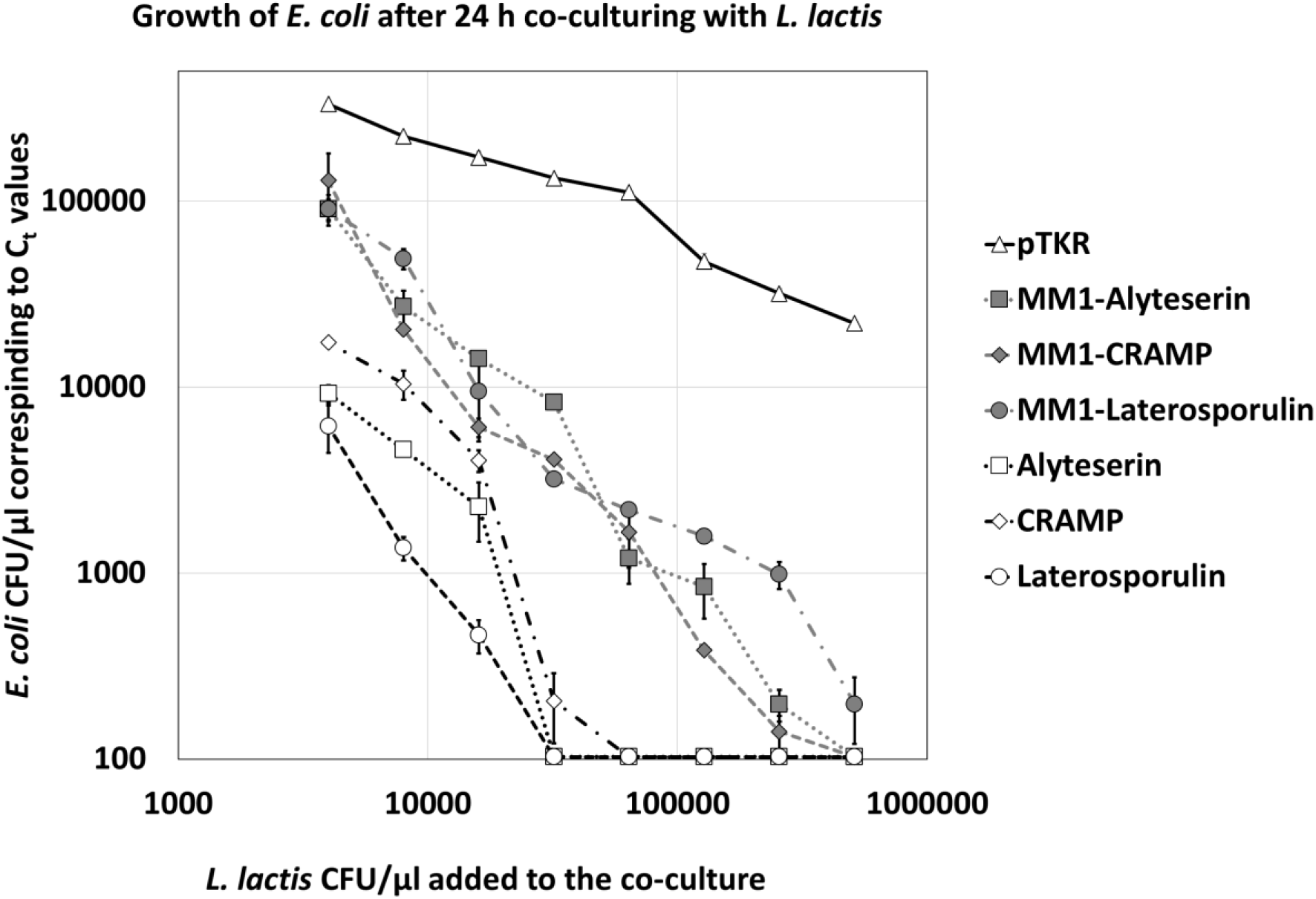
The CFU/ul of *E. coli* found in the co-culture after 24 hrs of incubation, estimated from the corresponding C_t_ values determined by qPCR, plotted against the CFU/ul of *L. lactis* added to the co-culture

### Chemically synthesized gAMP showed preferential bactericidal activity towards H. pylori over E. coli

Bacterial inhibition assay with the chemically synthesized peptide demonstrated that Alyteserin when conjugated with the MM1 fragment domain had preferential inhibitory action against *H. pylori* over the non-target bacteria *E. coli*. The MIC recorded against *H. pylori* and *E. coli* for both Alyteserin and MM1-Alyteserin is shown in Table 2 and the mean values are shown graphically in Figure 5. The MIC against *H. pylori* for Alyteserin and MM1-Alyteserin were not significantly different even though MM1-Alyteserin had a lower MIC but the MIC against *E. coli* the MIC for MM1-ALytserin was significantly higher (p<0.01) than that of Alyteserin, thus showing that MM1-Alyteserin shows significantly lower bactericidal activity towards the non-target *E. coli* compared to *H. pylori*. This echoes the results seen when the *L. lactis* cloned with AMPs and gAMPs were co-cultured with *H. pylori* and the non-target bacteria *E. coli* and *L. plantarum*.

**TABLE 1.** The peptide sequences of 3 AMPs and their corresponding gAMPs with Multimerin1 fragment fused to the N-terminus separated by a linker

| AMP/ gAMP | Peptide Sequence |
| --- | --- |
| Laterosporulin | ACQCPDAISGWTHTDYQCHGLENKMYRHVYAICMNGTQVYCRTEWGSSC |
| Multimerin1(MM1)-Laterosporulin | MQKMTDQVNYQAMKL TLLQKSGGGSACQCPDAISGWTHTDYQCHGLENKMYRHVYAICMNGTQVYCRTEWGSSC |
| Alyteserin | GLKDIFKAGLGSLVKGIAAHVAN |
| MM1-Alyteserin | MQKMTDQVNYQAMKL TLLQKSGGGSGLKDIFKAGLGSLVKGIAAHVAN |
| CRAMP | ISRLAGLLRKGGEKIGEKLKKIGQKIKNFFQKLVPQPE |
| MM1-CRAMP | MQKMTDQVNYQAMKL TLLQKSGGGSISRLAGLLRKGGEKIGEKLKKIGQKIKNFFQKLVPQPE |
| Blue = Multimerin1, Red = linker |  |

**TABLE 2.** Minimum Inhibitory Concentration (MIC) exhibited by chemically synthesized Alyteserin and MM1-Alyteserin against *H. pylori* and *E. coli*.

| Bacteria | MIC ($\mu\text{M}$) | Alyteserin | MM1-Alyteserin | P-value |
| --- | --- | --- | --- | --- |
| *H. pylori* | Mean | 83.8 | 61.2 | 0.1475 |
|  | SE | 5.429 | 10.842 |  |
| *E. coli* | Mean | 25.2 | 160.8 | 0.0004 |
|  | SE | 1.906 | 11.80 |  |

**FIGURE 5.**
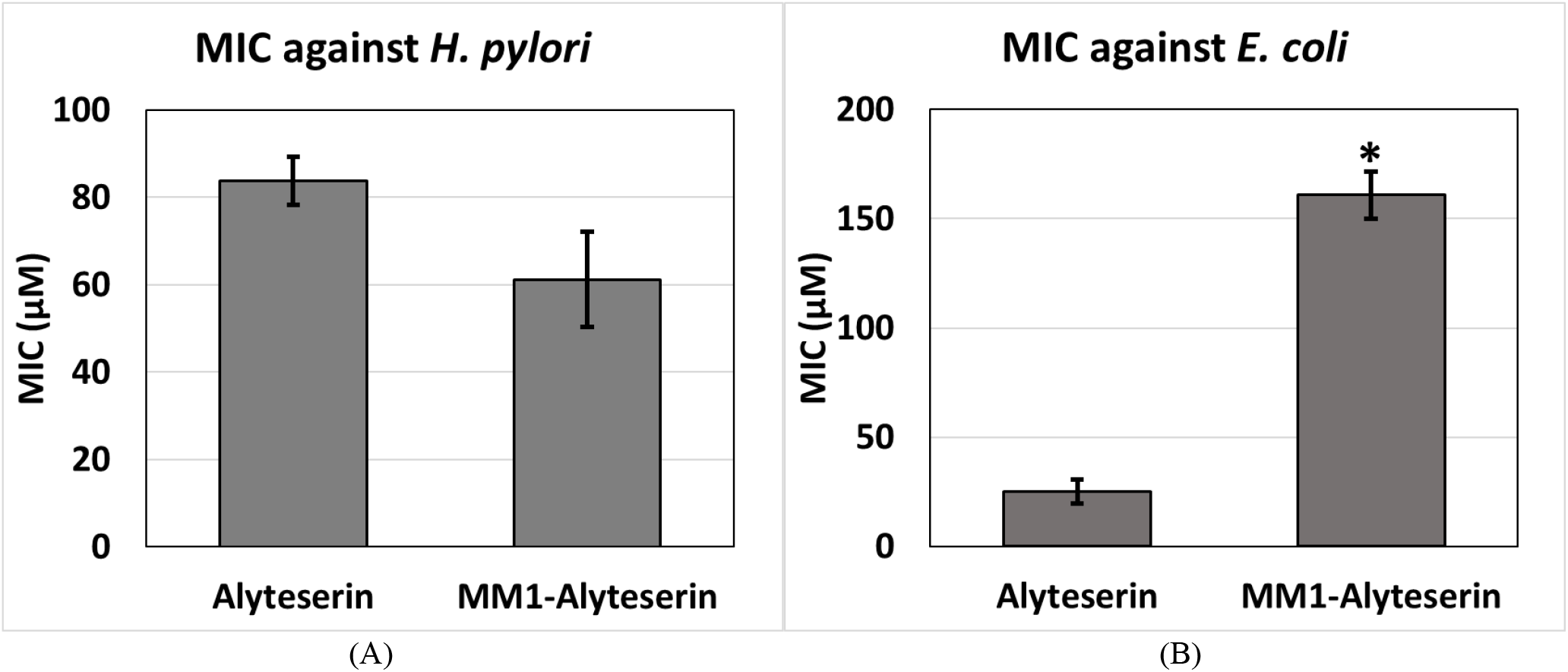
MIC values (µM) of Alyteserin and MM1-Alyteserin against *H*.*pylori* (A) and *E. coli* (B)

## Discussion

Both the *in vitro* assays using *L. lactis* cloned with AMP/gAMP and using chemically synthesized Alyteserin/MM1-Alyteserin demonstrated effective inhibtion of *H. pylori*. Co-culture assays showed that the *L. lactis* cloned with gAMPs that the Multimerin-1 fragment attached to it had similar efficacy against *H. pylori* to that of the *L. lactis* cloned with their AMP counterparts, with MM1-Alyteserin being the only gAMP that had significantly better microbicidal effect than its AMP counterpart. The measure of microbicidal effect was done by the estimating the CFU/µl of *H. pylori* in the co-culture after overnight incubation with a particular concentration of bioengineered *L. lactis*, the lesser *H. pyloti* load the better microbicidal action exhibited. So, we can assume that attachment of MM1 fragment did not necessarily turn the AMP a more potential killer of the target bacteria, *H. pylori*. But when looking at the effect on the other two non-*Helicobacter* species; *E. coli* and *L. plantarum*, we see that the gAMPs with MM1 fragment had a significant drop in their microbicidal effect as a higher CFU/ul of gAMP-*L. lactis* were required to get a similar drop in non-target bacterial load exhibited by a lower CFU/ml dose of AMP-*L. lactis*. Hence, similar concentration of gAMP-*L. lactis* that caused a significant reduction in *H. pylori* CFU/µl did not have the same effect on either *E. coli* or *L. plantarum* unlike the AMP-*L. lactis* which had similar microbicidal effect on all three bacteria. This shows that the gAMPs released by the *L. lactis* did not kill the non-target bacteria as it would *H. pylori*, proving that it has achieved an extent of specificity towards *H. pylori*. The fragment of MM1 that we used was found to have the highest dissociation coefficient (KD) with the vacA toxin on the surface of the *H. pylori* and may facilitate the attachment of the AMP to the bacteria. This enhanced attachment may not enhance the toxic effect that much as the amount of gAMP molecules approaching the bacterial cell membrane is not significantly different than the AMP molecules and attachment of gAMP molecules to the membrane above the saturation concentration does not necessarily make it more toxic. But the MM1 fragment may be an impairment to the attachment of the gAMP molecules to the non-target bacterial cell membrane and hence deceasing attachment which translates to decreased toxicity. This effect has also been seen in previous literature where targeting moieties did not necessarily enhance the toxic activity but rather decreased action against the other non-target bacteria. This ensures that the gAMPs have a certain specificity towards the bacteria they are designed to kill and sparing other non-target species which may be beneficial to the host microbiome.

When using a chemically synthesized version of the peptide Alyteserin and MM1-Alyteserin too reflected the preferential microbicidal tendency when their MICs were determined against *H. pylori* and *E coli*. Alyteserin is a class of AMPs among which Alyteserin-1 types show specific activity against Gram negative bacteria (Conlon et al., 2009). We used a specific sequence of an Alyteserin 1 peptide that has been used against Gram negative bacteria like *E. coli* (Conlon et al., 2009), *Salmonella* sp. (Volzing et al., 2013), *Acinetobacter* sp. (Conlon et al., 2010). The assay showed it to have similar MIC against *E. coli* as reported previously (∼25 µM) but that MIC was increased significantly when using MM1-Alyteserin whereas there was no significant changes between the MICs of either Alyteserin and MM1 Alyteserin against *H. pylori*, albeit the MM1 Alyteserin having a lower MIC against *H. pylori* though it was not significant. This again proves the validity of our argument for using MM1 as a targeting domain and corroborates the data we got with co-culture assay using cloned *L. lactis* to inhibit *H. pylori* instead of synthesized peptides as the peptides have a danger of getting degraded when administered orally. And since both the naked peptide and the cloned *L lactis* showed efficacy in killing *H. pylori* we can use the latter as a model of therapeutic against the pathogen instead of the former and making it a more suited candidate for oral administration.

